# Cytoplasmic aggregation of uranium in human dopaminergic cells after continuous exposure to soluble uranyl at non-cytotoxic concentrations

**DOI:** 10.1101/2020.07.16.205831

**Authors:** Asuncion Carmona, Francesco Porcaro, Andrea Somogyi, Stéphane Roudeau, Florelle Domart, Kadda Medjoubi, Michel Aubert, Hélène Isnard, Anthony Nonell, Anaïs Rincel, Eduardo Paredes, Claude Vidaud, Véronique Malard, Carole Bresson, Richard Ortega

## Abstract

Uranium exposure can lead to neurobehavioral alterations in particular of the monoaminergic system, even at non-cytotoxic concentrations. However, the mechanisms of uranium neurotoxicity after non-cytotoxic exposure are still poorly understood. In particular, imaging uranium in neurons at low intracellular concentration is still very challenging. We investigated uranium intracellular localization by means of synchrotron X-ray fluorescence imaging with high spatial resolution (< 300 nm) and high analytical sensitivity (< 1 μg.g^-1^ per 300 nm pixel). Neuron-like SH-SY5Y human cells differentiated into a dopaminergic phenotype were continuously exposed, for seven days, to a non-cytotoxic concentration (10 μM) of soluble natural uranyl. Cytoplasmic submicron uranium aggregates were observed accounting on average for 62% of the intracellular uranium content. In some aggregates, uranium and iron were co-localized suggesting common metabolic pathways between uranium and iron storage. Uranium aggregates contained no calcium or phosphorous indicating that detoxification mechanisms in neuron-like cells are different from those described in bone or kidney cells. Uranium intracellular distribution was compared to fluorescently labeled organelles (lysosomes, early and late endosomes) and to fetuin-A, a high affinity uranium-binding protein. A strict correlation could not be evidenced between uranium and the labelled organelles, or with vesicles containing fetuin-A. Our results indicate a new mechanism of uranium cytoplasmic aggregation after non-cytotoxic uranyl exposure that could be involved in neuronal defense through uranium sequestration into less reactive species. The remaining soluble fraction of uranium would be responsible for protein binding and the resulting neurotoxic effects.

## 1. Introduction

Natural uranium is a ubiquitous element in the environment, resulting in human population exposure to low but unavoidable concentrations from air, water and food (ATSDR, 2013). Natural uranium is a weakly radioactive element, its adverse health effects are mainly due to its chemical toxicity rather than its radiotoxicity (Keith *et al.*, 2008; ATSDR, 2013). Once ingested or inhaled, uranium circulates in the blood stream as soluble hexavalent uranium (uranyl) resulting in its accumulation in particular in bones and kidneys where severe acute toxic effects were documented (Keith *et al.*, 2008). High uranium exposures occur mostly in some occupational settings, typically for workers from uranium processing industries but also from phosphate fertilizer industry. Noteworthy, relatively high chronic exposure levels may also be found in geographical areas with elevated levels of naturally occurring uranium (Frisbie *et al.*, 2009; Frisbie *et al.*, 2013; Bjørklund *et al.*, 2017; Bjørklund *et al.*, 2020). Regardless the route of exposure, a small amount of uranium is able to reach the brain and may exert neurotoxic effects (Fitsanakis *et al.*, 2006; Houpert *et al.*, 2007; Dinocourt *et al.*, 2015). Rodent models exposed to uranium showed neurobehavioral changes such as increased locomotor activity, perturbation of the sleep-wake cycle, decreased memory, and increased anxiety. At the molecular level, such neurological alterations were caused by the disruption of the acetylcholine, serotonin, and dopamine neurotransmitter systems (Bussy *et al.*, 2006; Barber *et al.*, 2007; Dinocourt *et al.*, 2015). We recently reported the alteration of monoamine oxidase B expression in human dopaminergic cells exposed to natural uranium (Carmona *et al.*, 2018). The dopaminergic system was also impaired after chronic exposure of *C. elegans* to depleted uranium leading to the specific degeneration of dopaminergic neurons (Lu *et al.*, 2020). Further studies are needed to better understand the mechanisms of uranium-induced neurotoxicity on dopaminergic neurons. Such data are required to evaluate in particular whether long-term exposure to low uranium concentrations could affect neurological functions in humans.

Human SH-SY5Y cell line was selected as a study model. Human SH-SY5Y cells are frequently used as human *in vitro* neuronal model since these cells may acquire a neuronal dopaminergic phenotype after a differentiation procedure (Presgraves *et al.*, 2004). We selected a 7-day temporal window of continuous soluble uranyl exposure in order to approach *in vitro* the conditions inducing molecular effects reflecting more continuous rather than acute exposures. In these experimental conditions, we previously evidenced the intracellular isotopic fractionation of natural uranium, suggesting the existence of a high affinity transporter protein for uranyl uptake in SH-SY5Y dopaminergic cells (Paredes *et al.*, 2016). We also found a significant alteration of dopamine catabolic pathway beginning at non-cytotoxic (10 μM) uranyl continuous exposure (Carmona *et al.*, 2018). We addressed the question of uranium intracellular distribution using micro-PIXE (Particle Induced X-ray Emission) imaging, a technique applied with a spatial resolution of 2 μm and a limit of detection of about 10 μg.g^-1^ for uranium imaging. Micro-PIXE imaging revealed the uranium localization in cytoplasmic regions, particularly observable after continuous exposure to sub-cytotoxic (125 μM) and slightly toxic (250 μM) uranyl concentrations (Carmona *et al.*, 2018). The nature of these uranium-rich cytoplasmic regions and their existence for lower concentration exposures have yet to be determined.

The aim of the present study was to shed light on the mechanisms of uranium distribution in SH-SY5Y neuron-like cells after non cytotoxic continuous uranyl exposure. We assessed uranium subcellular distribution after continuous 7-day exposure to a non-cytotoxic uranyl concentration (10 μM) selected on the basis of our previously reported cytotoxicity results (Carmona *et al.*, 2018). We applied Synchrotron X-ray Fluorescence (SXRF) imaging, a technique with high spatial resolution (300 nm at Nanoscopium beamline, SOLEIL synchrotron) and high analytical element sensitivity (< 1 μg.g^-1^ per 300 nm pixel). We performed a cellular correlative microscopy approach by coupling SXRF imaging to epifluorescence microscopy for the identification of organelles and/or proteins fluorescently labeled (Roudeau *et al.*, 2014; Carmona *et al.*, 2019; Das *et al.*, 2019). Previous studies using transmission electron microscopy (TEM) indicated that after acute exposure to high uranium concentrations (> 300 μM) of alveolar macrophages, renal, or bone cells, uranium precipitates forming large needle-shape structures within cytoplasmic vesicles such as lysosomes (Hengé-Napoli *et al.*, 1996; Mirto *et al.*, 1999; Carrière *et al.*, 2008; Pierrefite-Carle *et al.*, 2017). We thus designed our study in order to evaluate, at lower non-cytotoxic concentration, the possible uranium co-localization with the cellular vesicular pathway (early endosomes, late endosomes, and lysosomes). In addition, we investigated the potential interaction of uranium with fetuin-A, a high affinity uranium-binding protein (Basset *et al.*, 2013).

## 2. Material and methods

### 2.1. Reagents and solutions

Eagle’s minimum essential medium (EMEM, 30-2003; ATCC), F12 medium (21765-029; Life Technologies), fetal bovine serum (FBS) (30-2020; ATCC), and penicillin/streptomycin (15070-063; Gibco-Thermo Fisher Scientific) solutions were used to prepare the culture medium for cell growth and exposure experiments. TrypLE Express 1×/EDTA (12605-010; Gibco-Thermo Fisher Scientific) was used for the trypsinization of cells. Phosphate buffered saline (PBS) (pH 7.4) free of CaCl2 and MgCl2 (10010-015; Gibco-Thermo Fisher Scientific) was used to wash the cells after trypsinization. Retinoic acid (RA) and 12-O-tetradecanoylphorbol-13-acetate (TPA) used for cell differentiation were purchased from Sigma-Aldrich (R2625 and P8139, respectively). According to our previous work (Carmona *et al.*, 2018), 3 mg.mL^-1^ RA solution was prepared in sterile dimethyl sulfoxide (DMSO, Sigma-Aldrich) under a nitrogen atmosphere in opaque tubes and stored at −80°C. TPA was re-suspended at 1 mg.mL^-1^ in sterile DMSO, and the solution was stored at −20°C. For organelles labeling we used: CellLight™ Early Endosomes-GFP BacMam 2.0 ThermoFisher Scientific C10586, CellLight™ Late Endosomes-RFP BacMam 2.0 ThermoFisher Scientific C10589, and CellLight^™^ Lysosomes-RFP BacMam 2.0 ThermoFisher Scientific C10589. Fetuin-A from fetal calf serum (F2379, Sigma) was first purified by size exclusion chromatography (TSKgel G3000-SW, Sigma-Aldrich) before being labeled with Alexa Fluor^®^ 488-NHS ester (Molecular Probes) using succimidyl ester chemistry. From their specific absorbance at 280 and 495 nm for the fetuin-A and the Alexa Fluor^®^ 488 dye, respectively, the concentration of the Alexa Fluor^®^ 488 labeled fetuin-A was 5.9 mg.mL^-1^ with 1.4 as final labeling degree.

### 2.2. Cells and dopaminergic differentiation

SH-SY5Y human neuroblastoma cells were purchased from ATCC USA (CRL-2266™, Lot 59740436). Cells were cultured according to ATCC guidelines. The complete growth medium consisted in a 1:1 ratio of EMEM (ATCC), and F12 Medium (Sigma-Aldrich), supplemented with 10% FBS (ATCC), 100 μg.mL^-1^ penicillin, and 100 μg.mL^-1^ streptomycin (Gibco-Thermo Fisher Scientific). Complete growth medium was renewed every 3 to 4 days. Cells were cultured at 37°C in air with 5% carbon dioxide. SH-SY5Y cells were terminally differentiated into dopaminergic neuron-like cells according to the protocol of Presgraves *et al.* (2004) and adapted to our experimental conditions (Carmona *et al.*, 2018). In brief, at day one, cells were seeded at a density of 25,000 cells.cm^-2^ onto sample holders adapted to SXRF and treated with 10 μM RA during 2.5 days to initiate the differentiation. Then the medium was renewed using fresh complete growth medium with 80 nM TPA and cells were maintained in culture for 3.5 more days. We demonstrated the validity of this protocol to induce a dopaminergic phenotype by Western blot analysis of tyrosine hydroxylase expression (Carmona *et al.*, 2018).

### 2.3. Organelles and fetuin-A fluorescent labeling

Early, late endosome and lysosome cellular organelles were labeled on living cells by means of the CellLight™ transitory transfection kits (ThermoFisher). These transfection kits provide insect virus baculovirus for the expression of fluorescent protein-signal peptide fusions in mammalian cells. Specific signal peptides are located exclusively in the selected organelles allowing highly specific organelle labeling. We checked that the fluorescence signal was stable up to 7 days after transfection enabling the transitory transfection before the 7-day period of uranium exposure (Fig. 1). In brief, after the RA-TPA differentiation period, and 16h prior to uranyl exposure, cells were incubated with fresh complete culture medium containing 15 μL of CellLight™ reagent per mL, corresponding to about 30 particles of baculovirus per cell in culture. Due to the different fluorescence colors selected (early and late endosome in green and lysosome in red), a double simultaneous transfection was also achievable. Cell exposure to fluorescently labeled Alexa Fluor^®^ 488 fetuin-A was carried out after uranium exposure (Fig. 1). Complete cell medium containing 36 μg.mL^-1^ of fluorescently labeled Alexa Fluor^®^ 488 fetuin-A was used to expose cells for 1.5 h before the live cell images acquisition. All cells, either labeled for organelles or for fetuin-A, were additionally labeled with a blue fluorescent nuclear dye, Hoechst 33342, just prior to fluorescent live imaging acquisition at 5-10 μg.mL^-1^ for 10 minutes at 37°C (Roudeau *et al.*, 2014; Carmona *et al.*, 2019; Das *et al.*, 2019). Live cell imaging of the fluorescent labels was recorded with an epifluorescence microscope (Olympus BX51).

**Fig. 1.**
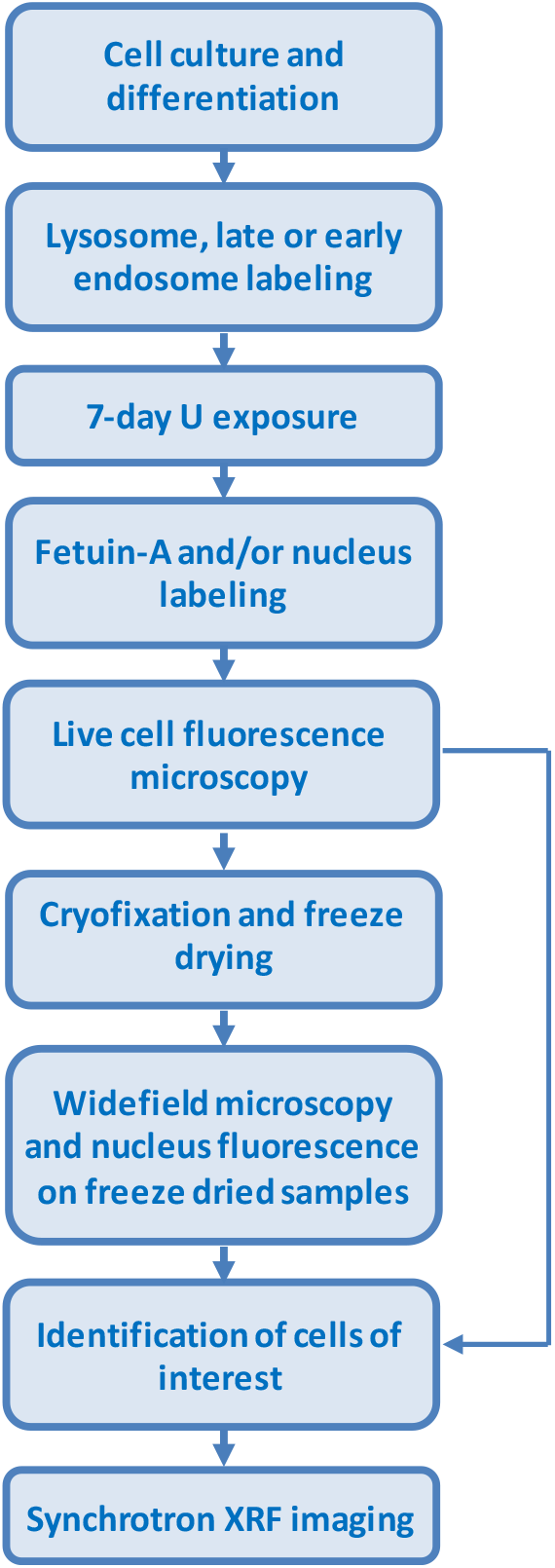
Schematic workflow for SH-SY5Y cell culture, organelle labeling, 7-day continuous exposure to natural uranyl, live cell microscopy and cryo-processing before SXRF imaging.

### 2.4. Preparation of the uranium exposure solution

A in-house (Laboratoire de développement Analytique, Nucléaire, Isotopique et Elémentaire (LANIE), DES, CEA Saclay) natural uranium stock solution at concentration of 151 mM was obtained by dissolving natural uranium oxide powder U_3_O_8_ in 0.5 M ultrapure HNO3 (SCP Science). This solution was diluted into a buffer, made of 0.1 mol.L^-1^ NaHCO_3_ (Sigma-Aldrich), 0.1 mol.L^-1^ Na_2_CO_3_ (Sigma-Aldrich), 0.15 mol.L^-1^ NaCl (Sigma-Aldrich) and 0.05 mol.L^-1^ Tris (Biorad) in ultrapure water (Fisher), in order to obtain an intermediate solution of soluble uranyl (the hexavalent cation UO_2_^2+^ is spontaneously formed in aqueous buffers and in presence of oxygen), with a pH compatible with cell culture experiments (pH 8-8.5). In order to avoid uranyl precipitation, the uranium stock solution was added dropwise into the buffer, with a dilution 1:5 (w/w) ratio to reach 30 mM uranium concentration. Thus, uranium was stabilized in this intermediate solution under the tris-carbonate uranyl (VI) complexes [UO_2_(CO_3_)3]^4-^, the most common soluble uranium species found in intracellular, interstitial, and plasma fluids (Sutton & Burastero, 2004). Uranium concentration in the intermediate solution was daily measured by quadrupole Inductively Coupled Plasma Mass Spectrometry (ICP-MS, X7 Series; Thermo Fisher Scientific) for one week showing that the intermediate solution was stable, without any uranium precipitate, at least during 7 days. The intermediate uranyl solution was prepared the day before use for cell exposure and stored at room temperature. The 10 μM uranyl exposure solution was prepared in complete cell culture medium by dilution of the 30 mM intermediate natural uranium solution. Uranium content after dilution in the culture medium was measured systematically by ICP-MS to verify that the exposure solution contained the expected uranium concentration. A comprehensive study of uranium speciation in a 10 μM solution of culture medium, prepared using this protocol, was previously performed and demonstrated the full solubility of uranium and the absence of precipitation in the exposure solution (Paredes *et al.*, 2016, Paredes *et al.*, 2018).

### 2.5. Cell culture and exposure to natural uranyl

Sample holders for SXRF consist in a thin (2 μm) ultrapure polycarbonate film stretched over a polyether ether ketone (PEEK) frame with a 5 mm x 5 mm window where cells are grown (Perrin *et al.*, 2015). Sample holders were sterilized in analytical grade ethanol, rinsed with ultrapure water and dried in sterile conditions. To enhance cell attachment, polycarbonate films were coated with a solution of 0.01 mg.mL^-1^ of bovine serum albumin, 0.01 mg.mL^-1^ of human fibronectin, and 0.03 mg.mL^-1^ of collagen type I (all products from Sigma-Aldrich) prepared in complete growth medium. Before cell seeding onto polycarbonate films (day 0), SH-SY5Y cells were first cultured in plastic flasks for 10 days after thawing following standard methods as described above. At day 0, SH-SY5Y cells were harvested, counted, and 25,000 cells.cm^-2^ seeded onto each SXRF sample holder. Cells were thereafter grown on polycarbonate films, differentiated with RA-TPA during 7 days, transfected for the selected organelle labeling and finally exposed to defined concentrations of natural uranyl for 7 more days (Fig. 1). Uranyl exposure solutions at 10 μM were prepared in complete cell growth medium by dilution of the 30 mM intermediate uranyl solution as detailed in the previous section.

### 2.6. Sample preparation for SXRF experiments

The protocol for sample preparation was adapted from optimized methods for element imaging in single cells using SXRF and micro-PIXE (Perrin *et al.*, 2015; Carmona *et al.*, 2019; Das *et al.*, 2019). Immediately after live cell microscopy, cells were rinsed with a 150 mM ammonium acetate buffer solution, pH 7.4, prepared in ultrapure water (Fisher) to remove traces of extracellular elements. Then cells were cryofixed by rapid plunge freezing in isopentane chilled with liquid nitrogen down to −160°C, and freeze-dried at −90°C under primary vacuum (Christ Alpha 1-4 freeze-drier). After 3 days of freeze drying, samples were slowly brought back to room temperature and ambient pressure. A second epifluorescence microscopy run on freeze dried samples was carried out in order to check for sample preservation and to record the positions of the selected fluorescent cells (Fig. 1). The stability of Hoechst 33342 nuclear dye fluorescence after freeze drying allows the localization of the nuclear areas on freeze-dried cells (Roudeau *et al.*, 2014; Das *et al.*, 2019). Finally, samples were stored into a desiccator box until SXRF analysis. Before the start of the experiments, the samples were mounted in a homemade double confinement sample holder to ensure a proper safe storage during the measurements.

### 2.7. SXRF analyses and data processing

SXRF analyses were carried out at Nanoscopium beamline (Somogyi *et al.*, 2015) from the French national facility, synchrotron SOLEIL. SXRF element distributions were acquired by using a 17.6 keV X-ray beam just above the L3 absorption edge of uranium. The incoming X-ray beam was focused to 300 x 300 nm^2^ size with a Kirkpatrick-Baez nano-focusing mirror-pair, resulting in about 10^10^ ph.s^-1^ flux. In order to map the uranium distributions in single cells, around 2-4 hours per cell was required with 300 ms dwell time per pixel and 300 nm pixel size. A home-made MATLAB code was used for on-line data treatment at the Nanoscopium beamline for extracting the corrected elemental distribution maps from the identified X-ray region of interests. 3D XRF data cubes were also extracted by the MATLAB code for further data-processing by PyMca software. In a second step, SXRF elemental distribution images were extracted from fitted XRF spectra using the PyMca software in order to take into account potential spectral overlapping (Sole *et al.*, 2007). Individually and for 18 different analyzed cells, mean SXRF spectra corresponding to both the whole cell and to the regions showing uranium aggregates were extracted and fitted using PyMca software. This permitted to obtain the number of counts in the uranium Lα peaks corresponding to the entire cell and the aggregates, respectively. Image compositions were performed using Fiji open source package (http://imagej.net) (Schindelin *et al.*, 2012). Fiji was also used to measure the number and size of uranium aggregates. The GraphPad Prism 8 software was used for graphical purposes (http://www.graphpad.com/).

## 3. Results and discussion

### 3.1. Uranium distribution

In our previous work (Carmona *et al.*, 2018), we determined the cytotoxicity thresholds of soluble hexavalent uranium (uranyl) on SH-SY5Y differentiated cells for a continuous 7-day exposure. We found that exposure to 250 μM uranyl resulted in 15% of cell viability inhibition, while exposure to 125 μM was considered the limit below which no cytotoxic effects were observable. We selected a concentration of 10 μM, far from the cytotoxicity threshold, to study neurotoxic effects of uranyl at non-cytotoxic concentrations and found a decrease in monoamine oxidase B (MAO-B) expression. In the present study we addressed the issue of uranium distribution in SH-SY5Y differentiated cells exposed for 7 days continuously to this non-cytotoxic uranyl concentration (10 μM). SXRF imaging revealed that uranium is actually detected in the cells but not homogeneously distributed as it could have been expected after cell exposure to soluble uranyl. Uranium is localized in the cytoplasm within uranium-rich structures as illustrated in Fig. 2 for three representative examples. We could evaluate the size of these uranium rich-areas and found that, in the same cell, they vary from about 300 nm (1 pixel) to around 1.5 μm diameter (5 pixels) (Fig. 2). It is noteworthy that single uranium-rich structures smaller than 300 nm will appear with an apparent size of 300 nm on the SXRF images. The size of the smallest uranium-rich structures is very likely below 300 nm. These cytoplasmic uranium-rich structures will be further referred as uranium aggregates, because of their high mass per unit area, in opposition to the other intracellular fraction of uranium that will be further referred as soluble uranium. Uranium aggregates of submicron size were systematically found in the cytoplasm of all SH-SY5Y cells imaged by SXRF and never detected outside the cells (Fig. 2 and 4–7).

**Fig. 2.**
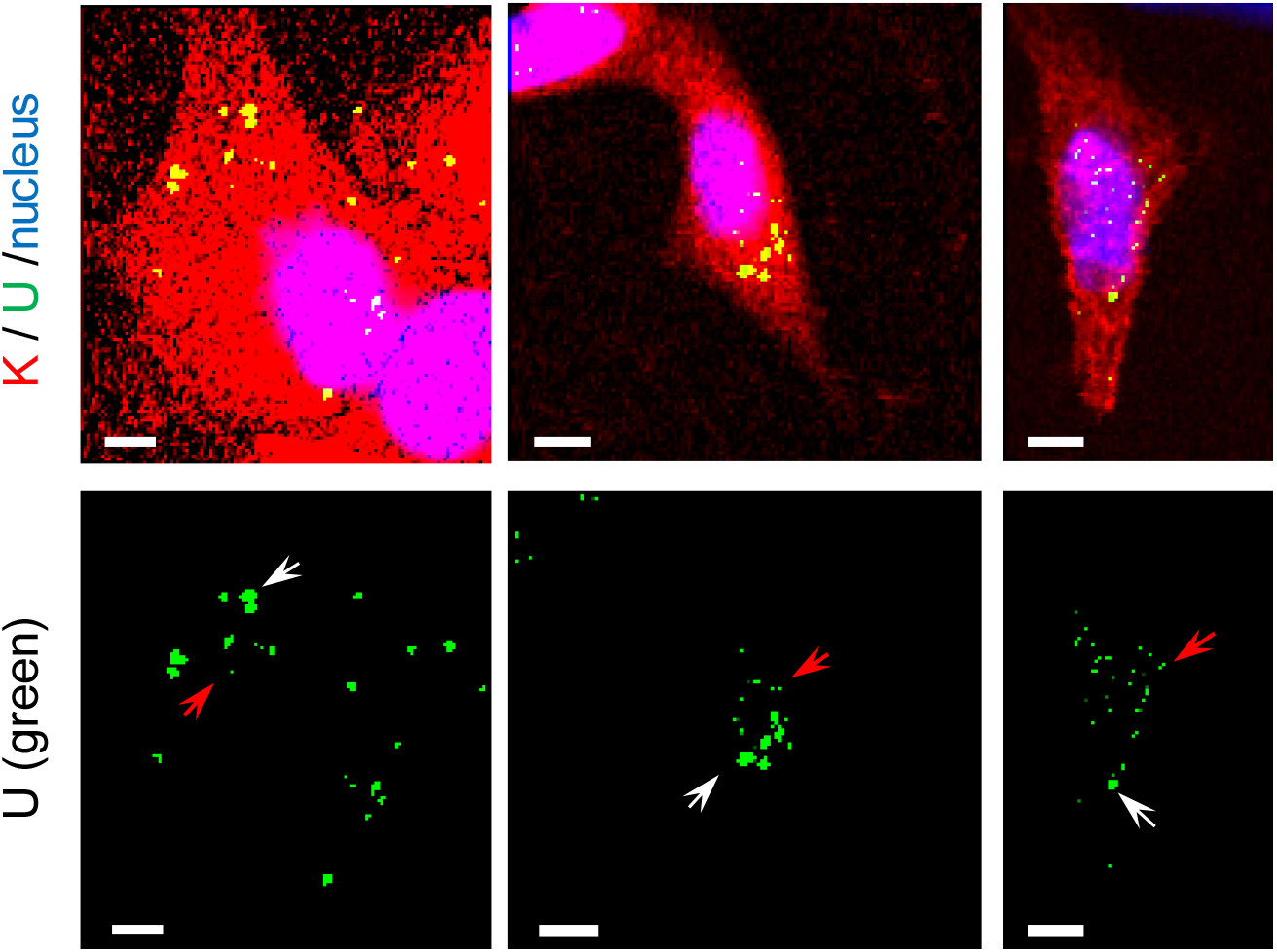
Representative examples of uranium distribution in SH-SY5Y cells exposed to 10 μM uranyl. The first row shows the merged images of SXRF elemental maps of potassium (red) and uranium (green) together with the nucleus fluorescence labeling (blue). The second row shows the uranium distribution alone (green) to evaluate the size of the aggregates. White arrows indicate some large (0.9-1.5 μm) uranium aggregates and red arrows smaller ones (<300 nm). Scale bars: 5 μm.

**Fig. 3.**
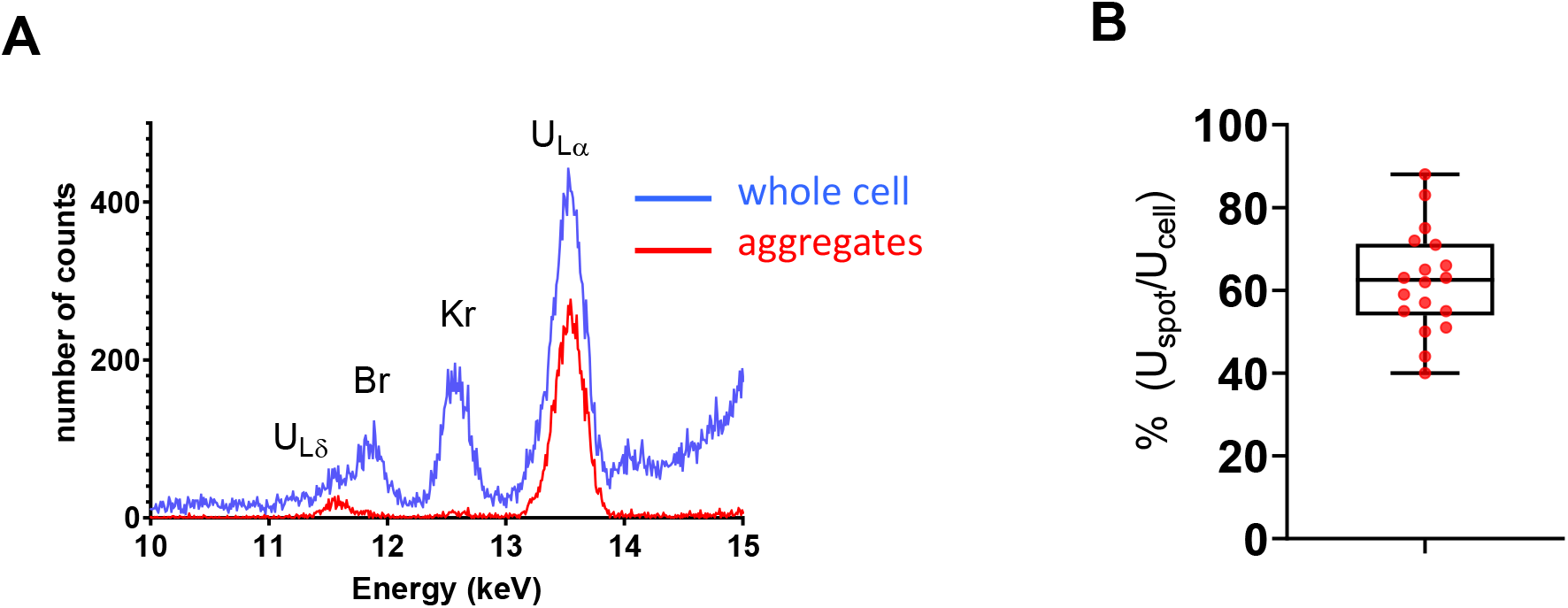
Quantification of uranium content in aggregates versus the whole cell. A) Representative SXRF spectra obtained from the selection of a whole cell (blue spectrum) and from the uranium aggregates within the same cell (red spectrum). SXRF analyses are performed in air explaining the presence of Kr. B) Box and whiskers plot of uranium aggregates (U_spot_) versus total cellular uranium (U_cell_), expressed as a percentage and calculated for 18 different cells.

**Fig. 4.**
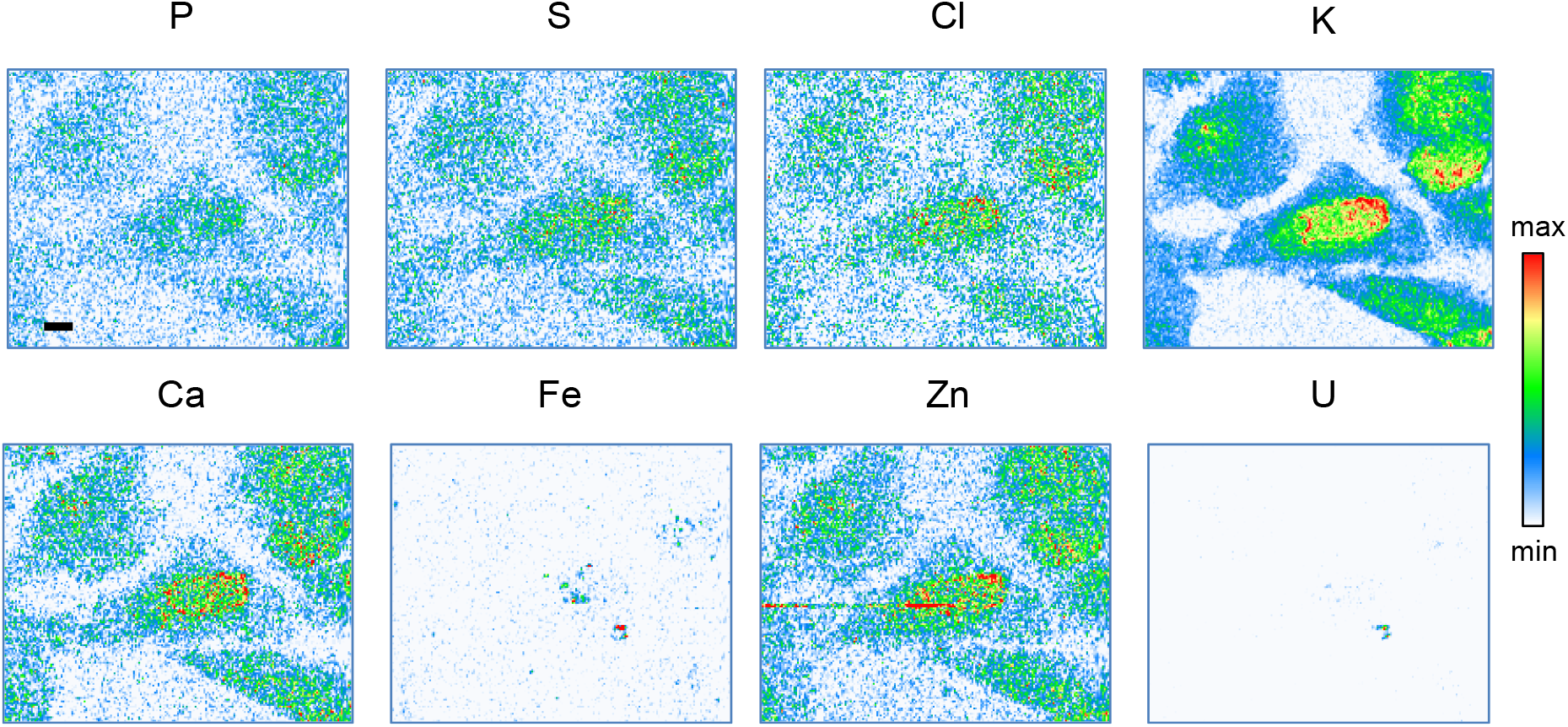
SXRF multi-elemental imaging. Representative example of P, S, Cl, K, Ca, Fe, Zn and U distributions in SH-SY5Y cells exposed to 10 μM uranyl solution for 7 days. Except for Fe and U, the other detected elements display a homogeneous intracellular distribution with X-ray fluorescence signal proportional to the cell thickness. Scale bar: 5 μm.

**Fig. 5.**
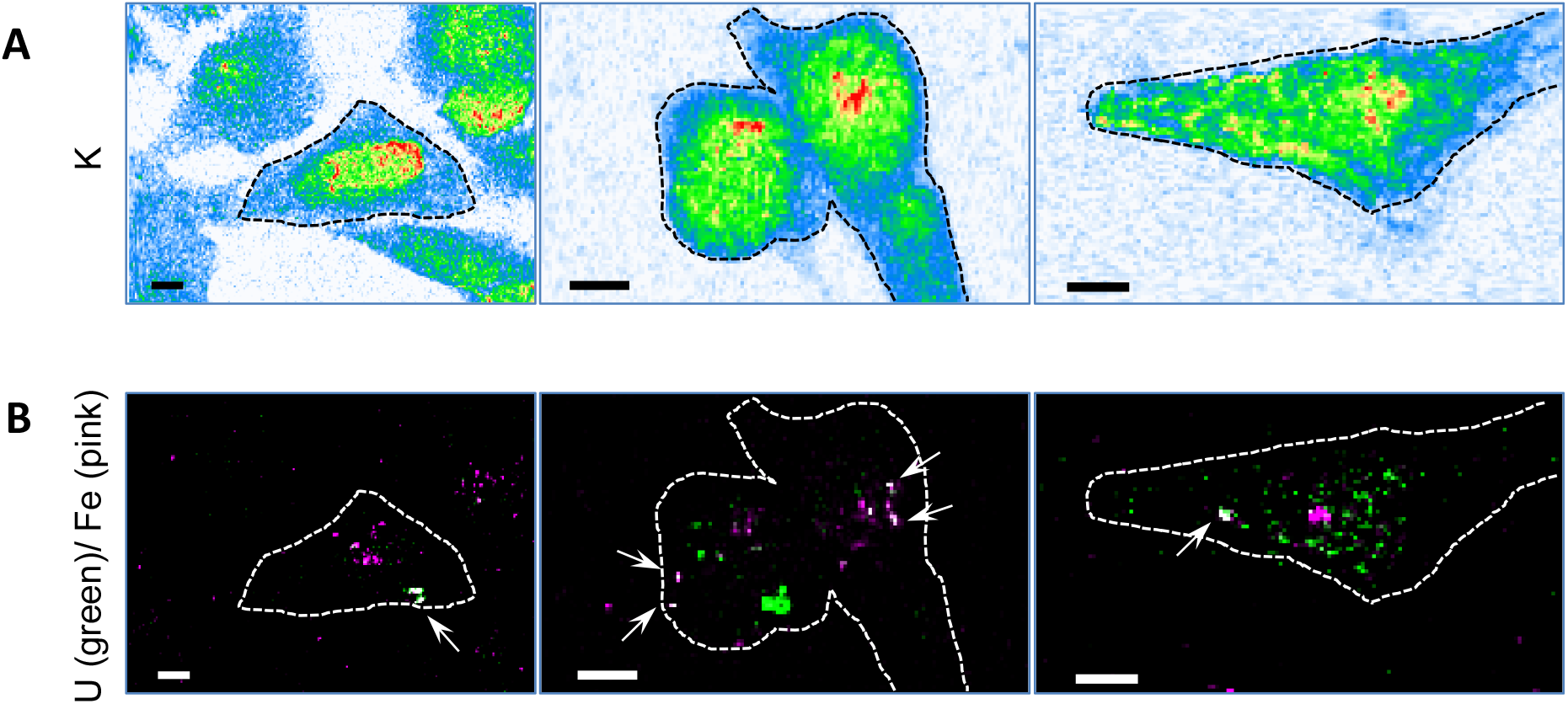
Uranium and iron partial co-localization in SH5Y-SY cells. A) The first row shows the potassium elemental maps where the cell boundaries are drawn with a black dotted line. B) The second row shows the merged images of uranium (green) and iron (magenta) and the cell boundaries (white dotted line). Co-localization of iron together with uranium is revealed by the white color of the merged images and highlighted by white arrows. Scale bars: 5 μm.

**Fig. 6.**
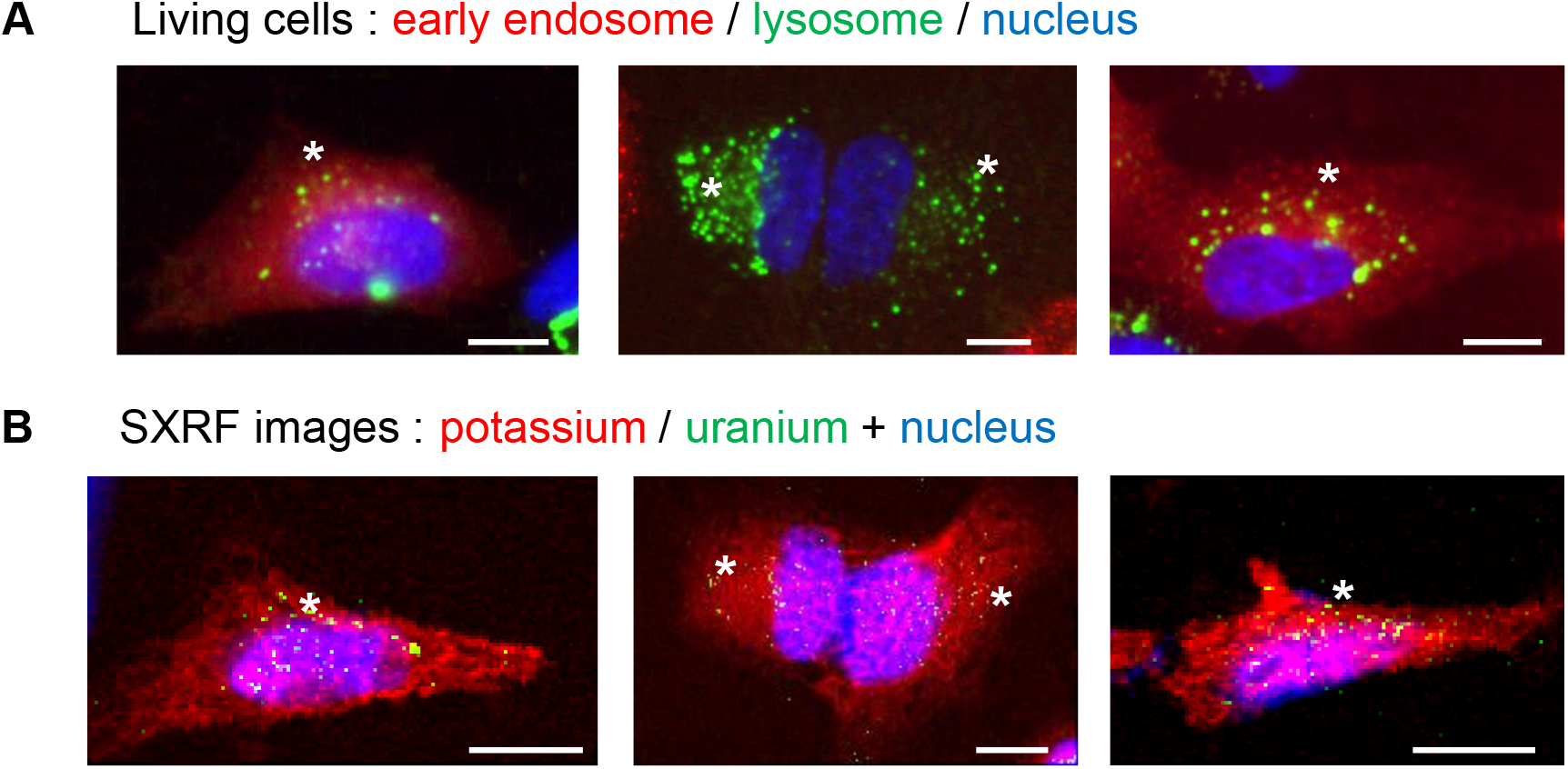
Uranium distribution and comparison with nucleus, lysosome and early endosome localization. A) Live cell microscopy fluorescence imaging of early endosome (red), lysosome (green) and nucleus (blue). B) SXRF imaging of potassium (red), uranium (green) and fluorescence imaging of nucleus (blue) after freeze-drying of the cells presented in A). White asterisks indicate the presence of uranium aggregates in cytoplasmic areas where lysosome fluorescence is dense. Scale bars: 10 μm.

**Fig. 7.**
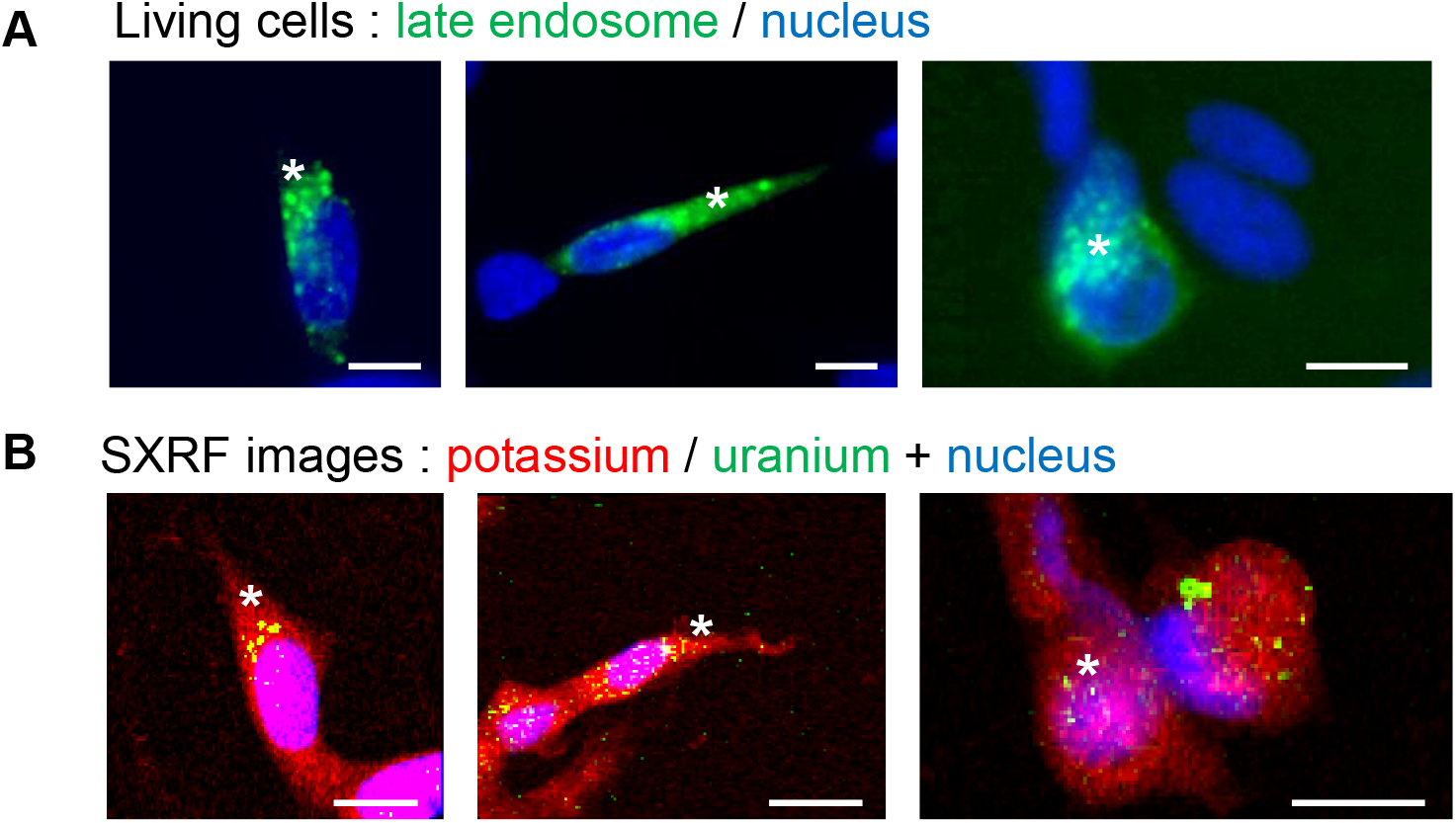
Uranium distribution and comparison with late endosome localization. A) Live cell microscopy fluorescence imaging of late endosome (green) and nucleus (blue). B) SXRF imaging of potassium (red), uranium (green) and fluorescence imaging of nucleus (blue) after freeze-drying of the cells presented in A). White asterisks indicate the presence of uranium aggregates in cytoplasmic areas where late endosome fluorescence is dense. Scale bars: 10 μm.

To go further in the analysis of uranium distribution in cells, we quantified the ratio of uranium content within the submicron-aggregates *vs* the total uranium content in the whole cells (Fig. 3). As illustrated for one representative example in figure 3A, X-ray fluorescence indicates the presence of uranium, characterized by its ULα emission peak at 13.6 keV, within the aggregates (red spectrum) but also shows a larger signal from the whole cell (blue spectrum). We determined the total number of counts corresponding to the uranium peaks in both cases, inside the aggregates and for the whole cell, for 18 different cells. We found that, on average 62 ± 13% (mean ± SD, n=18) of uranium is in the aggregate form, and this ratio varies between 40% and 88% from cell to cell (Fig. 3B). These results indicate that uranium is present within cells in two forms, soluble and aggregated, with about two thirds of the total cellular uranium trapped in the aggregate state.

At high exposure concentrations, typically above 300 μM, uranium can be easily imaged in cells by TEM due to the formation of electron-dense aggregates. Such structures were reported after 24h exposure to 800 μM uranium in LLC-PK1 kidney cells (Mirto *et al.*, 1999), after 12h and 24h exposure to 300 μM uranium in NRK-52E kidney cells (Carrière *et al.*, 2008), and after 24h exposure to 300 μM uranium in UMR-106 osteosarcoma cells (Pierrefite-Carle *et al.*, 2017). Uranium aggregates are not seen by TEM for lower exposure concentrations, such as for example at 100 μM in osteosarcoma cells. The exposure time before observing the uranium aggregates is an important parameter. Although visible after 24h in osteosarcoma cells exposed to 300 μM uranium, these aggregates are not observed after 6h exposure at the same concentration (Pierrefite-Carle *et al.*, 2017). The size of the uranium aggregates increases with time as observed in NRK-52E kidney cells (Carrière *et al.*, 2008). Using more sensitive element imaging techniques, such as secondary ion mass spectrometry (SIMS), uranium aggregates were observed in HepG2 liver cancer cells and in IMR32 neuroblastoma cells after 24h exposure to 100 μM depleted uranium nitrate, but not below this concentration (i.e. 10 and 50 μM) (Rouas *et al.*, 2010). Our results indicate the existence of cytoplasmic uranium aggregates in cells exposed to non-cytotoxic concentrations of soluble uranyl, i.e. 10 μM. The 7-day continuous exposure to uranyl solution might be a favorable factor to allow the formation of uranium aggregates. In addition, by using SXRF, a very sensitive element imaging technique, we were able not only to detect uranium in cellular aggregates at low concentrations, but also to evidence the presence of uranium in cells outside the aggregates. Using SXRF with a 1 μm beam size, uranium was also detected in cytoplasmic regions of kidney cells from a rat model of uranium-induced acute renal toxicity (Homma-Takeda *et al.*, 2015). The cytoplasmic preferential localization of uranium was described as well in human kidney cells exposed for 1h to 500 μM uranium using uranyl-specific fluorescent antibodies and confocal microscopy (Reisser-Rubrecht *et al.*, 2008).

The physico-chemical form of uranium, and in particular its solubility, is a key parameter to address its toxicity (ATSDR, 2013; Brown *et al.*, 2014). A more soluble compound will exert a higher toxicity due higher bioavailability and higher diffusion in the body fluids and cells. The intracellular solubilization of metallic particles is a known mechanism of toxicity resulting in the specific toxicity of the soluble part of metallic species released in the cytosol (Ortega *et al.*, 2014). Conversely, the aggregation of soluble uranium species in SH-SY5Y cells would result in a protective effect by sequestering the diffusible uranyl species into less reactive uranium species. This mode of cell protection has been proposed to explain the decreased genotoxicity observed concomitantly with the intracellular precipitation of uranium in embryonic zebra-fish cells exposed to 250 μM uranyl solution (Pereira *et al.*, 2012). The mechanism of biologically driven uranium aggregation may explain the low cytotoxicity of uranium in SH-SY5Y cells. It is noteworthy that for a non-cytotoxic exposure condition, most of the intracellular uranium content (>60%) is found within submicron-aggregates. This proportion is likely to be higher considering that uranium may be present in submicron-aggregates of less than 300 nm size, the limit of identification imposed by the spatial resolution of SXRF. That being said, a non-negligible proportion of uranium remains in a soluble form within SH-SY5Y cells explaining the binding to intracellular proteins (Vidaud *et al.*, 2019) and the effects on monoamine enzymes (Carmona *et al.*, 2018) occurring at non-cytotoxic uranyl exposure concentration (10 μM).

### 3.2. Multi-elemental imaging

Hard X-ray SXRF is a multi-elemental imaging technique that enables the simultaneous detection of intracellular elements with atomic numbers > 13 (aluminum), through the detection of K- and L-lines X-ray emission. In addition to uranium, the other intracellular elements detected were phosphorous, sulfur, chlorine, potassium, calcium, iron and zinc, as illustrated in Fig. 4. This figure shows that only uranium and iron present a dotted distribution. The other elements are quite homogenously distributed, with an X-ray fluorescence intensity signal proportional to the cellular thickness, i.e. stronger in the nuclear region (Carmona *et al.*, 2008).

Early electron X-ray microanalysis studies indicated the presence of phosphorous in the uranium precipitates as observed in alveolar macrophages from rats exposed to uranium aerosols (Hengé-Napoli *et al.*, 1996), or in kidney cells exposed *in vitro* to 800 μM uranyl, both suggesting the precipitation of uranyl-phosphate (Mirto *et al.*, 1999). The speciation analysis of uranium using extended X-ray fine structure spectroscopy (EXAFS) in kidney cells exposed to 300 μM uranyl indicated the presence of both uranium phosphate and uranium carbonate (Carrière *et al.*, 2008). In osteosarcoma cells exposed to 300 μM uranyl, precipitates were identified as meta-autunite Ca[(UO2)(PO4)]2(H2O)11 by EXAFS (Pierrefite-Carle *et al.*, 2017). Phosphorous and uranium were co-localized in renal proximal tubules from rats exposed to uranyl acetate as shown by micro-PIXE and micro-EXAFS (Homma-Takeda *et al.*, 2019). In neuron-like cells exposed to 10 μM uranyl we did not observe phosphorous or calcium co-localization with uranium suggesting a different mechanism of aggregation. The metabolism of SH-SY5Y cells is expected to differ from that of osteoblasts, macrophages or kidney cells. For instance, Boonrungsiman *et al.* (2012) demonstrated the role of intracellular calcium phosphate in osteoblasts in the osteoblast-mediated apatite formation in bones. Our results suggest that, in neuron-like cells, uranium would not precipitate as calcium-phosphate or phosphate related compounds. Another uranium precipitate form identified by EXAFS is uranium carbonate in kidney cells (Carrière *et al.*, 2008). Since SXRF is not sensitive to carbon or oxygen detection, we cannot conclude about the actual presence of uranium carbonate in the observed aggregates. The speciation of uranium aggregates in SH-SY5Y cells still remains to be determined. Such characterization may constitute a very challenging analytical issue that would need to overcome the methodological limitations of micro-EXAFS since this technique requires a good knowledge of the expected uranium species to allow their identification (Porcaro *et al.*, 2018).

From our data, the only element that is partially co-localized with uranium is iron. Both elements display a granular cytoplasmic distribution. As illustrated in Fig. 5 for three representative examples of cells, merging iron and uranium distributions highlights that these elements are co-localized in some aggregates (see arrows in Fig. 5), although most of iron and uranium-rich areas in cells are not co-localized. It is noteworthy that iron homeostasis is modified following uranium exposure. For example, dense granular inclusions containing iron have been observed in kidney cells of rats chronically exposed to uranium, although uranium could not be detected in the iron-rich inclusions using electron X-ray microanalysis (Donnadieu-Claraz *et al.*, 2007). In addition, we have recently reported the increase of Fe/Mg ratio in SH-SY5Y cells after 7-day continuous exposure to 10 μM uranyl (Paredes *et al.*, 2018). Now, our result suggests that uranium might interact with iron storage pathways in SH-SY5Y cells. Interestingly, the ferritin genes, a cytosolic iron-storage protein, are highly up-regulated in SH-SY5Y cells exposed to cadmium, a neurotoxic metal (Forcella *et al.*, 2020). Although no data have been reported yet on the interaction of uranium with ferritin in human cells or tissues, it is known that uranium can bind to ferritin when human apoferritin solutions are loaded with uranyl (Hainfeld, 1992; Michon *et al.*, 2010). Our result might also be related to the identification of ferritin as one of the main uranium-binding protein in hepatopancreas cells in a model organism of environmental uranium-bioaccumulation (*Procambarus clarkii*) (Xu *et al.*, 2014).

Another candidate protein to bind iron and uranium in dopaminergic cells is alpha-synuclein. Alpha-synuclein is a metal-binding protein with high iron-binding affinity (Peng *et al.*, 2010). In rat dopaminergic neurons exposed to an excess of iron, the overexpression of alpha-synuclein leads to the increase of intracellular iron and its redistribution in the cytoplasm, within perinuclear α-synuclein inclusions (Ortega *et al.*, 2016). Recently, alpha-synuclein aggregation has been observed in dopaminergic cells from *C. elegans* chronically exposed to uranium, leading to the specific neurodegeneration of dopaminergic neurons in these worms (Lu *et al.*, 2020). Although the interaction of uranium with alpha-synuclein in human dopaminergic cells remains hypothetical, such eventuality would have significant consequences in understanding uranium neurotoxicity.

### 3.3. Uranium, organelles and fetuin-A imaging

In our previous work, we found the uranium localization in sub-cytoplasmic regions of differentiated SH-SY5Y human cells after 7-day continuous exposure to 10, 125 and 250 μM soluble uranyl (Carmona *et al.*, 2018). This result was obtained by means of a correlative imaging approach using micro-PIXE element imaging and optical fluorescence microscopy of nuclear dyes (Roudeau *et al.*, 2014). This sub-cytoplasmic uranium distribution suggested the potential accumulation of uranium in organelles. In this same study, we further compared uranium distribution to Golgi apparatus and lysosome fluorescence on cells exposed to 125 and 250 μM soluble uranyl. Uranium distribution was clearly different from the Golgi apparatus localization. Uranium and lysosomes had more similar distributions. Although micro-PIXE is characterized by good resolution (2 μm) and sensitivity (about 10 μg.g^-1^ for uranium), the uranium content was very close to the detection limit when the cells were exposed to 10 μM uranyl (Carmona *et al.*, 2018). This is why we now investigated uranium distribution by means of SXRF a more sensitive element imaging method (< 1 μg.g^-1^), and with higher spatial resolution (300 nm).

In order to identify the nature of the sub-cytoplasmic uranium distribution in SH-SY5Y cells, several organelles were labeled. We performed a triple fluorescent labelling of the nucleus, lysosome and early endosome using respectively the blue nuclear dye, Hoechst 33342, the red fluorescence CellLight™ Lysosomes-RFP and the green fluorescence CellLight™ Early Endosomes-GFP. Representative examples of intracellular uranium distribution with their relative optical fluorescent images are reported in figure 6. Our results show that the position of the uranium aggregates differs from the lysosome and early endosome localization. Although a strict correlation between uranium aggregates and the lysosomes is not observed, uranium aggregates are often localized in cytoplasmic areas where the fluorescence intensity of the lysosomes is dense (marked by the asterisks in fig. 6). Since the cell position and size may slightly vary between the epifluorescence microscopy and the SXRF images due to the freeze-drying procedure (Roudeau *et al.*, 2014), it cannot be completely excluded that some uranium aggregates could be related to lysosomes.

The nucleus and the late endosome were labeled using the Hoechst 33342 dye and the fluorescence CellLight^™^ Late endosome-GFP. Some representative examples of cells analyzed by SXRF are illustrated in figure 7. A strict correlation between uranium and the late endosome localization is not evidenced. However, similarly to what we described for the lysosome, uranium aggregates are found in the cellular regions of higher late endosome vesicles density (marked by the asterisks in Fig. 7), a partial co-localization cannot be completely excluded.

Uranium aggregates have been observed in lysosomes of alveolar macrophages from rats exposed to uranium nitrate aerosols (Hengé-Napoli *et al.*, 1996). Alveolar macrophages contain a high number of lysosomes and high activity of acid phosphatase, an intralysosomal enzyme, resulting in uranyl-phosphate precipitation. In osteosarcoma cells, uranium precipitate was identified in intracellular vesicles, such as lysosomes, multi-vesicular bodies or auto-phagosomes (Pierrefite-Carle *et al.*, 2017). As previously mentioned in the multi-elemental discussion section, the metabolism of SH-SY5Y neuron-like cells largely differs from that of osteoblasts, or from alveolar macrophages, explaining why uranium is not predominantly found in lysosomal or endosomal vesicles.

In the last epifluorescence labeling experiment we imaged Alexa-488 labeled fetuin-A and Hoechst 33342 nuclear dye. Some representative examples for comparison of uranium and fetuin-A distributions are displayed in figure 8. Fetuin-A is a serum glycoprotein and member of the cystatin protein family, mainly produced in the liver. Fetuin-A has been identified as the carrier protein in human serum showing the highest affinity for uranium-binding and among the main potential proteins responsible for uranium uptake (Basset *et al.*, 2013). Fetuin-A is expressed in the mature brain and may have a neuroprotective function in mature inflamed and ischemic human brains (Heinen *et al.*, 2018). In SH-SY5Y cells we found that Alexa-488 labeled fetuin-A is internalized within cytoplasmic vesicles (Fig. 8A), similarly to the endocytosis of fluorescently labeled fetuin-A already described for other cells such as vascular smooth muscle cells (Reynolds *et al.*, 2005; Chen *et al.*, 2007), or murine macrophages (Wang *et al.*, 2011). We investigated the hypothesis whether uranium could be co-localized with vesicles containing fetuin-A in SH-SY5Y cells. We could not identify a clear correlation between the optical fluorescence of fetuin-A and uranium X-ray emission. This result suggests that uranium and fetuin-A might be internalized via different routes.

**Fig. 8.**
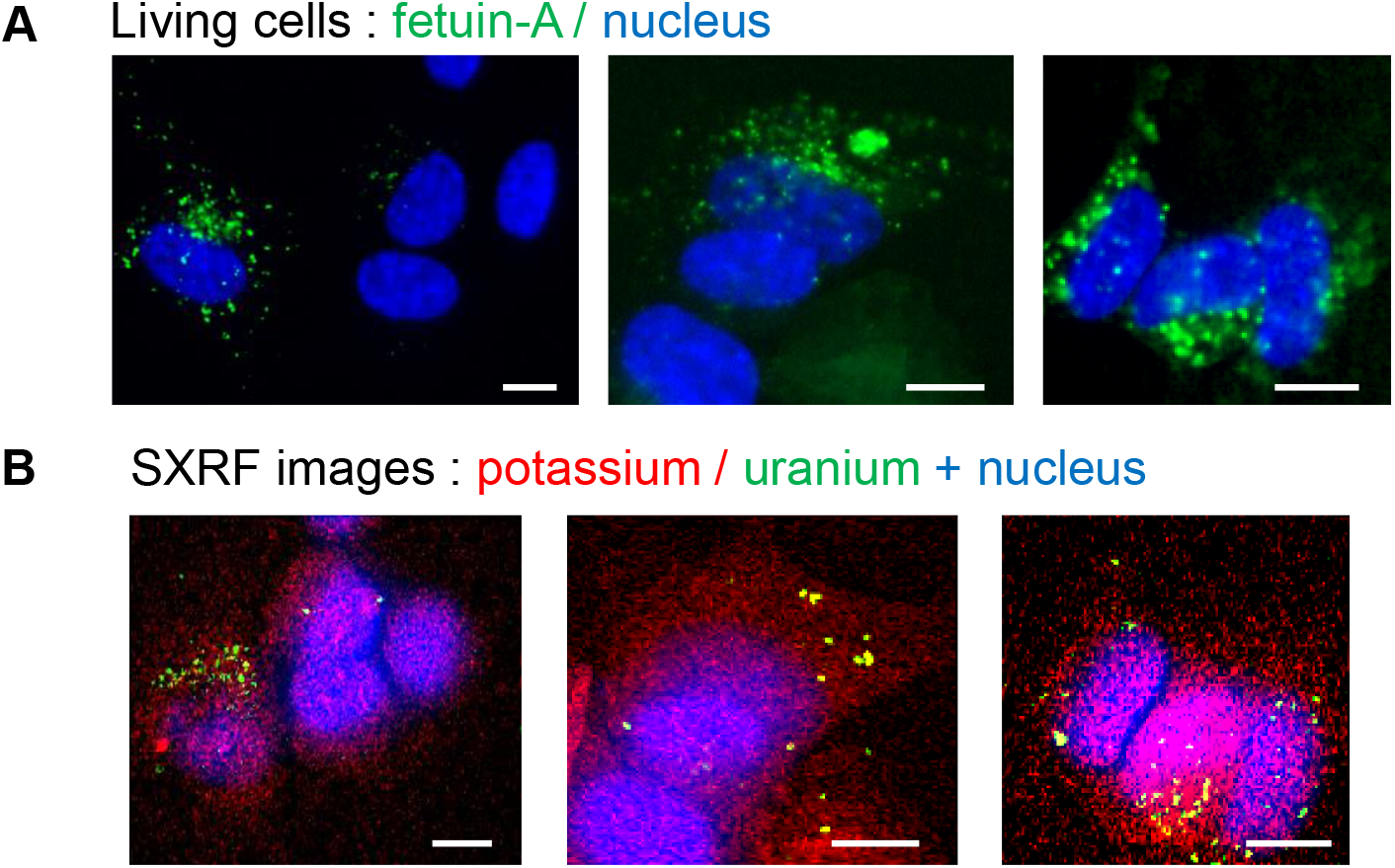
Uranium distribution and comparison with fetuin-A localization. A) Live cell microscopy fluorescence imaging of Alexa 488 labeled fetuin-A (green) and nucleus (blue). B) SXRF imaging of potassium (red), uranium (green) and fluorescence imaging of nucleus (blue) after freeze-drying of the cells presented in A). Scale bars: 10 μm.

## 4. Conclusions

In the last years, the uranium toxicology field has focused on the impact of uranium on the nervous system (Dinocourt *et al.*, 2015). However, the complex interaction between uranium and the human nervous system is far from being elucidated. In this context, we investigated the uranium intracellular distribution in neuron-like cells differentiated in a dopaminergic phenotype. In a previous study we revealed the alteration of the monoamine degradation pathway after uranium continuous exposure at non-cytotoxic concentration (Carmona *et al.*, 2018). We also mapped uranium distribution using PIXE chemical imaging with micrometric resolution and reported the presence of uranium in sub-cytoplasmic regions. In order to address the question of a possible uranium accumulation in specific cytoplasmic organelles we increased uranium element sensitivity and resolution using SXRF imaging. Furthermore, by means of selective optical fluorescence labeling, we could stain some potential organelles or proteins of interest in order to find possible correlations with uranium signal.

Thanks to the excellent sensitivity and spatial resolution of the SXRF technique, we could expose neuronal-like cells to 10 μM uranyl solutions and reveal the formation of uranium submicron-aggregates in the cytoplasm that could not be observed before with other techniques. The exact nature of these uranium aggregates and their mechanism of formation in SH-SY5Y cells remain elusive. Contrary to what could be described for higher exposure conditions, and in other cell types, phosphorous or calcium were not detected in the uranium submicron-aggregates. The mechanism of uranium cytoplasmic aggregation in neuron-like cells is therefore probably different from the mechanisms described in osteoblasts, which involve a calcium phosphate precipitation through biomineralization pathways, or in macrophages resulting in uranium precipitation in lysosomes due to the specific activity of lysosomal acid phosphatase in these cells.

In addition, we evidenced the co-localization of iron and uranium in some, but not all, uranium aggregates. This result suggests a potential interaction of uranium with intracellular iron storage proteins such as ferritin, a cytosolic protein which was demonstrated to sequester uranyl in biochemical assays. However, because only a small fraction of uranium aggregates contained iron, a direct interaction of uranium with iron storage protein may not account for the full mechanism of uranium aggregation. This result however indicates new avenues of research to elucidate the nature of uranium aggregates that may share common metabolic pathways with iron.

We also explored other hypotheses to identify the nature of uranium aggregates by comparing the distribution of uranium with vesicular organelles (lysosomes, early and late endosomes) and with vesicles containing fetuin-A. Our results do not suggest a strict co-localization of uranium with the endo-lysosomal pathway or with vesicles containing labeled fetuin-A. Therefore, considering the size of the uranium aggregates (<300 nm), the fact that they are not co-localized with phosphorous or calcium, and that they are not strictly co-localized with the endo-lysosomal and fetuin-A vesicles, it is likely that uranium aggregates could be formed directly within the cytosol of SH-SY5Y neuron-like cells.

We propose that uranium aggregation in SH-SY5Y neuron-like cells could be a mechanism of cell defense against uranyl neurotoxicity, starting at low levels of uranyl continuous exposure, resulting in the sequestration of uranium into less reactive species (i.e. ferritin-binding). How the soluble uranyl is progressively clustered into uranium dense cytoplasmic areas is still puzzling and will require further investigations. Clues might come from the coordination chemistry of uranyl with biological ligands, the chemical environment seen by uranium affecting its binding and aggregation. The magnitude of non-specific uranium–protein interactions is expected to be important in cells and may lead to the formation of multinuclear uranium clusters with proteins (Van Horn & Huang, 2006). Multinuclear uranium clusters with proteins could represent the early phase of formation of uranium cytosolic aggregates. Although we suggest that this mechanism could be protective, uranium aggregation with target neurotoxic proteins (i.e. alpha-synuclein) might contribute to neurotoxicity. Finally, it is important to highlight that a non-negligible proportion of uranium remains soluble in the cell. This soluble uranyl pool may account for the neurotoxic effects, such as the alteration of the monoaminergic metabolism, observed at non-cytotoxic uranyl exposure concentrations.

## Credit authorship contribution statement

**Asuncion Carmona:** Conceptualization, Methodology, Formal analysis, Investigation, Writing-Original Draft, Writing-Review & Editing, Visualization**. Francesco Porcaro:** Methodology, Formal analysis, Investigation, Writing-Original Draft**. Andrea Somogyi:** Resources, Investigation, Writing-Review & Editing**. Stéphane Roudeau:** Investigation, Writing-Review & Editing**. Florelle Domart:** Investigation**. Kadda Medjoubi:** Resources, Investigation. **Michel Aubert:** Investigation**. Hélène Isnard:** Investigation**. Anthony Nonell:** Investigation**. Anaïs Rincel:** Investigation**. Eduardo Paredes:** Conceptualization, Writing-Review & Editing**. Véronique Malard:** Conceptualization, Writing-Review & Editing, Funding acquisition**. Claude Vidaud:** Conceptualization, Resources, Writing-Review & Editing**. Carole Bresson:** Conceptualization, Investigation, Writing-Review & Editing, Funding acquisition**. Richard Ortega:** Conceptualization, Methodology, Investigation, Writing-Original Draft, Writing-Review & Editing, Supervision, Project administration, Funding acquisition.

## Acknowledgements

We acknowledge SOLEIL for provision of synchrotron radiation facilities and we would like to thank NANOSCOPIUM staff for assistance in using the beamline. The authors are grateful to Dr. Marta Garcia Cortes for the characterization of the Alexa Fluor^®^ 488 labeled fetuin-A solution.

